# Evidence for novel transient cell clusters in the neonatal mouse retina

**DOI:** 10.1101/2019.12.27.888792

**Authors:** Jean de Montigny, Vidhyasankar Krishnamoorthy, Fernando Rozenblit, Tim Gollisch, Evelyne Sernagor

**Author notes:** Equal contribution to the study (Vidhyasankar Krishnamoorthy, Fernando Rozenblit).

## Abstract

Waves of spontaneous activity sweep across the neonatal mouse retinal ganglion cell (RGC) layer, driven by directly interconnected cholinergic starburst amacrine cells from postnatal day (P) 0-10, followed by waves driven by glutamatergic bipolar cells. We found transient clusters of auto-fluorescent cells in the RGC layer during the period of cholinergic waves. They appear around the optic nerve head at P2 and become gradually displaced towards the periphery between P2-8 and then they disappear. Pan-retinal multielectrode array recordings reveal that cholinergic wave origins follow a similar, non-random developmental center-to-periphery pattern. Electrical imaging unmasks hotspots of dipole electrical activity occurring in the vicinity of wave origins. We propose that these activity hotspots are the sites for wave initiation and may originate from the transient cell clusters, reminiscent of activity in transient subplate neurons in the developing cortex, suggesting a universal hyper-excitability mechanism in developing CNS networks during the critical period for brain wiring.

## Introduction

During development, neural wiring is refined through activity-dependent processes (Blankenship and Feller, 2010; Luhmann et al, 2016). Spontaneous activity emerges long before sensory experience is possible, displaying unique expression patterns in different CNS areas. In the visual system, this activity is manifested by waves of spikes spreading across the retinal ganglion cell (RGC) layer (Meister et al, 1991). Several studies have demonstrated that retinal waves guide the development of visual connectivity (Huberman et al, 2008; Assali et al, 2014).

The cellular mechanisms underlying wave generation change with development, indicated by profound changes in the wave spatiotemporal features (Maccione et al, 2014). In mouse, the drive for wave generation/propagation switches from gap junction communication (Stage-1, prenatal) to cholinergic neurotransmission originating in starburst amacrine cells (SACs) (Stage-2, late gestation to P9) (Feller et al, 1996). Control then switches to glutamatergic bipolar cells before waves disappear around eye opening (Stage-3, P10-13). During Stage-2, SACs make direct homotypic connections, leading to lateral activity propagation across their network (Zheng et al, 2004). Experimental (Zheng et al, 2006; Ford et al, 2012) and theoretical studies (Butts et al, 1999; Hennig et al, 2009; Matzakos-Karvouniari et al, 2019) suggest that SACs play a fundamental role in defining wave dynamics by driving both wave initiation and propagation. Active SACs impose a refractory period, creating boundaries for activity propagation and controlling wave frequency. However, wave properties are not static during the prolonged Stage-2 period, exhibiting a gradual increase in wave frequency and size, followed by substantial shrinkage from P7 (Maccione et al, 2014). This suggests that Stage-2 wave initiation and propagation mechanisms may be more complex than originally assumed. Here we report for the first time that transient clusters of cells are present in the RGC layer during Stage-2 waves, and we propose that these cells may act as wave pacemakers.

## Results

While exploring the pan-retinal expression of cholinergic cells during Stage-2 waves (P2-10) using immunostaining for the acetylcholine synthesizing enzyme Choline Acetyl Transferase (ChAT), we made the serendipitous discovery of sparse cellular clusters (Figure 1, green dots) in the RGC layer (marked with RGC-specific RBPMS ((Kwong et al, 2010; Rodriguez et al., 2014), red). They form an annulus of tight clusters around the optic nerve head (ONH) at P2. As development progresses, the annulus expands, reaching the retinal periphery around P6-7. The clusters then begin to disintegrate and completely disappear by P10, coinciding with the switch from Stage-2 to Stage-3 waves.

**Figure 1:**
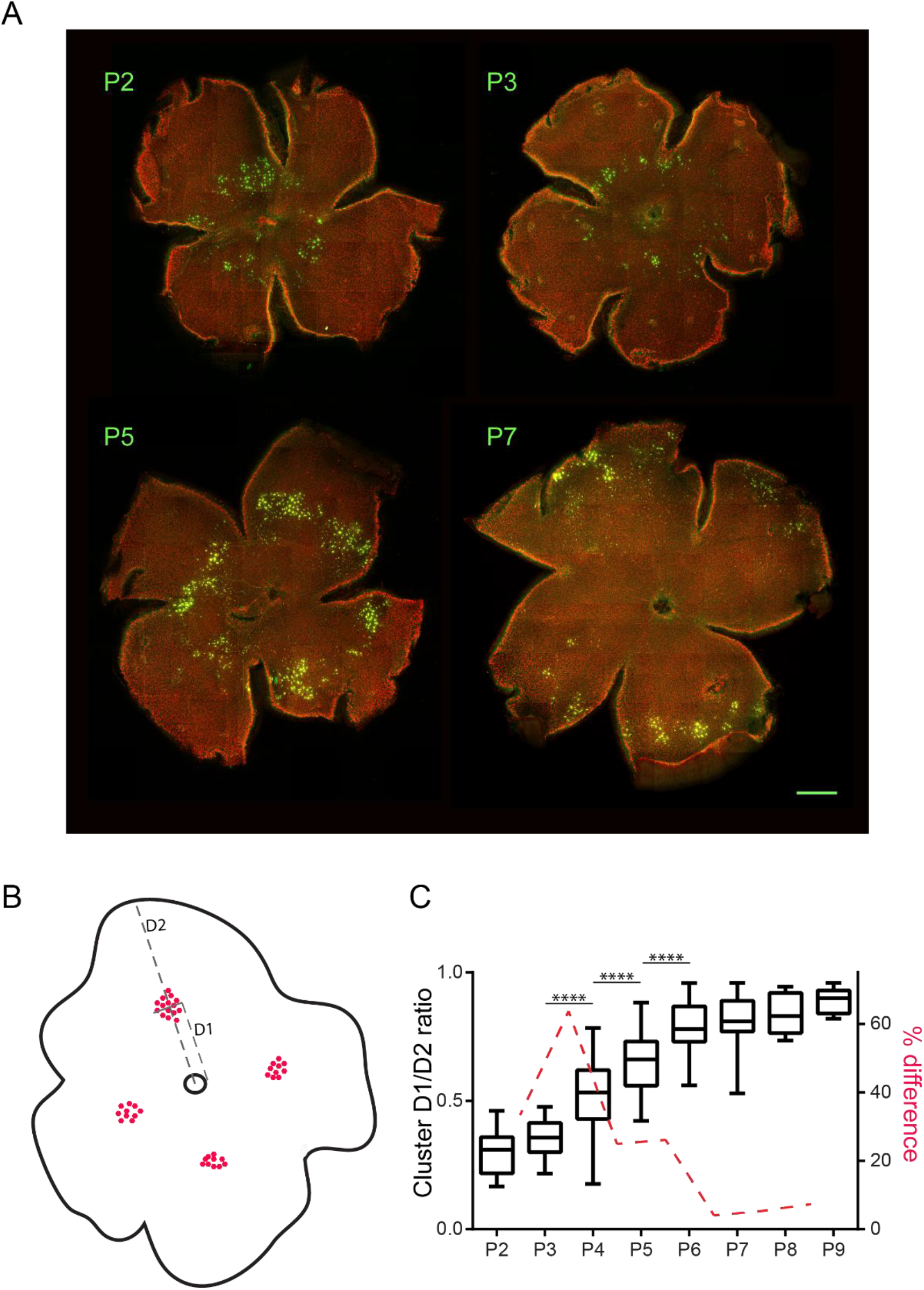
Retinal cell clusters expansion from center to periphery during the first postnatal week. **A**: Mouse retinal wholemounts stained for ChAT (green) and RBPMS (red). Scale bar: 500 μm. **B**: Method for calculating the relative position of clusters between the ONH (small black circle in the middle) and periphery. Cluster cells are represented by red dots. D1: distance from center of ONH to center of cluster. D2: distance between center of cluster to periphery. **C**: Box plot showing developmental changes in D1/D2 ratio. Each box illustrates the median (horizontal line) and interquartile range, with minimum and maximum values (whiskers). Asterisks indicate significant changes between consecutive days (One-way ANOVA with post-hoc Tukey test). The red dotted line illustrates the percentage difference in values between consecutive days, showing peak difference between P3 and P4 and no further changes from P6 onwards.

We quantified the clusters’ eccentricity during development (Figure 1B,C). The most substantial expansion occurs between P3-5, with no further eccentric movement beyond P6.

These cells auto-fluoresce across a broad wavelength spectrum (Figure 2A), and therefore their identity cannot be determined using immunofluorescence. They are larger than ACs and even than RGCs and they are present only in the RGC layer, but not in the inner nuclear layer (Figure 2B). Retinal sections stained for vesicular acetylcholine transporter (VAChT) to visualize SACs and the typical sublaminas generated by their processes in the inner plexiform layer reveal that the cluster cells are in close physical proximity to SACs and may even contact them (Figure 2C, arrow).

**Figure 2:**
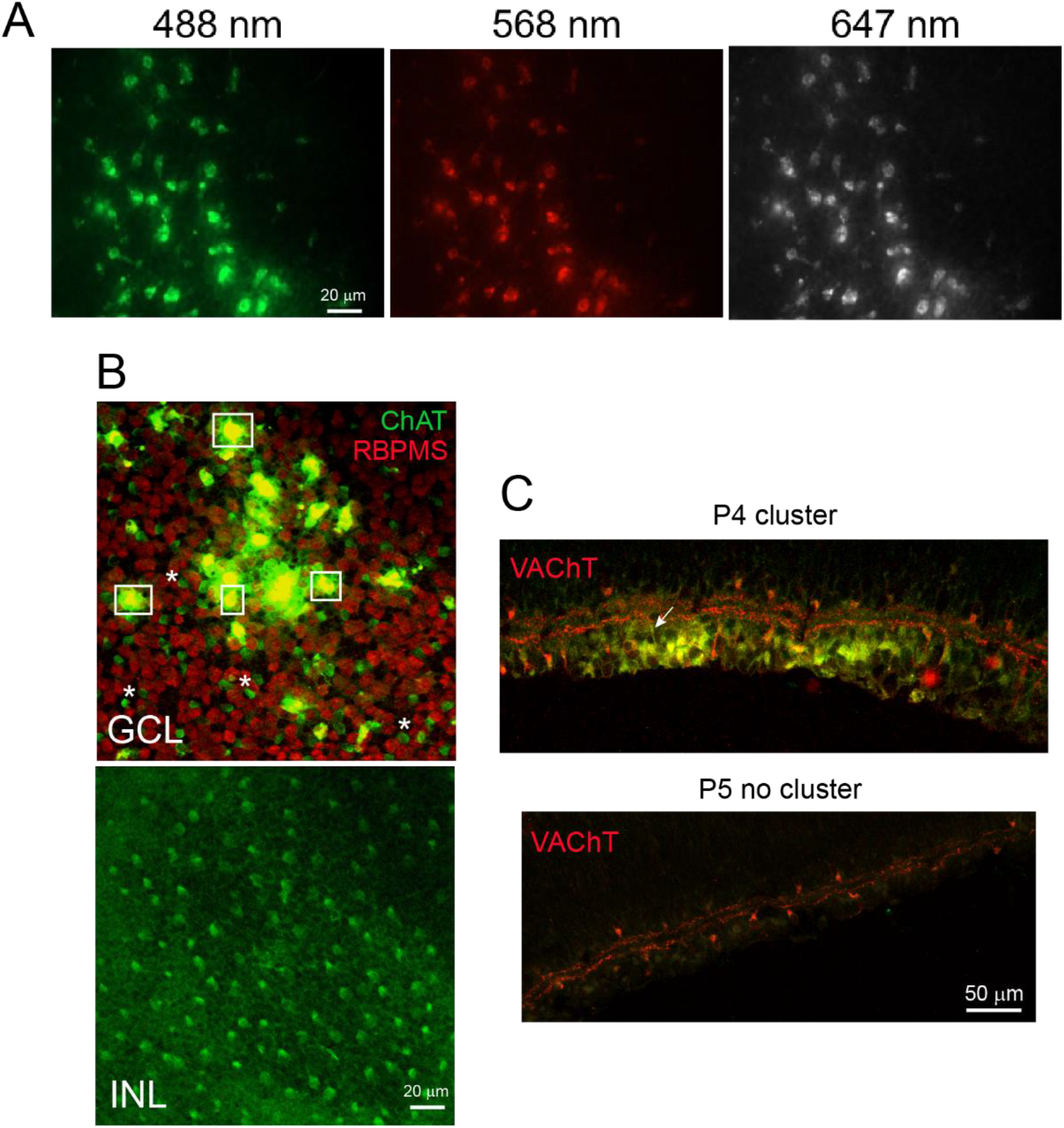
Cluster cells are auto-fluorescent and reside in the ganglion cell layer. **A**: one cluster (P5 retina) visualized at three different wavelengths without any immunolabelling, indicating intrinsic fluorescent signals. **B**: P5 cluster viewed at the ganglion cell layer (GCL) level and at the inner nuclear layer (INL) level. Green: ChAT; red: RBPMS. The cluster cells are visible only at the GCL level (examples in white boxes) and they are much larger than SACs (white asterisks) or RGCs (in red). At the INL level, there are only SACs. **C**: Retinal sections showing VAChT (red) expression in a cluster area (P4 retina) and in an area devoid of clusters (P5 retina). SACs express VAChT, exhibiting the typical double laminar expression in the IPL flanked by cell bodies in the INL and GC. In areas devoid of clusters, only the SAC expression pattern can be seen.

The cell clusters are present only during the period of Stage-2 waves, expanding from center to periphery between P2-7. We have previously shown that wave sizes also increase from P2-6 (Maccione et al., 2014). These coincidental events made us wonder whether these clusters might be involved in wave generation. If so, we predict that wave origins would also migrate in an eccentric fashion during the first postnatal week. To explore this possibility, we recorded waves between P2-13 using large-scale multielectrode arrays (MEAs) with 4,096 active electrodes spanning an area of 5.12×5.12 mm^2^, electrode pitch 81 μm, large enough to cover the entire retinal surface at all ages (Figure 3). Wave origins (determined as the xy coordinates of the initial wave center of activity trajectory, see Maccione et al., 2014) were aligned with the image of the retina itself (Figure 3A, red dots, and B, green dots) and then classified as either central or peripheral (Figure 3A and 3C, see Methods). Figure 3D shows how the periphery/center ratios of wave origins change with development. There is a very clear centrifugal shift, (with ratio values reaching >1) occurring after P3, with maximum change between P3-5, exactly like for the cluster locations (Figure 1C). That trend disappears with the emergence of the late Stage 3 glutamatergic waves. At that point, the periphery-to-center ratio drops to the lowest values, corroborating our previous findings that glutamatergic waves are small activity hotspots that tile the entire retina (Maccione et al., 2014). When wave origins are randomized between C and P (Monte Carlo randomization, 10 repetitions), the trend disappears and ratio values approach 1 for all developmental stages (Figure 3D, red). In stark contrast with all previous studies on retinal wave dynamics, our findings thus suggest, that during Stage-2, wave initiation is highly non-random, following the same centrifugal trajectory as the cluster cells.

**Figure 3:**
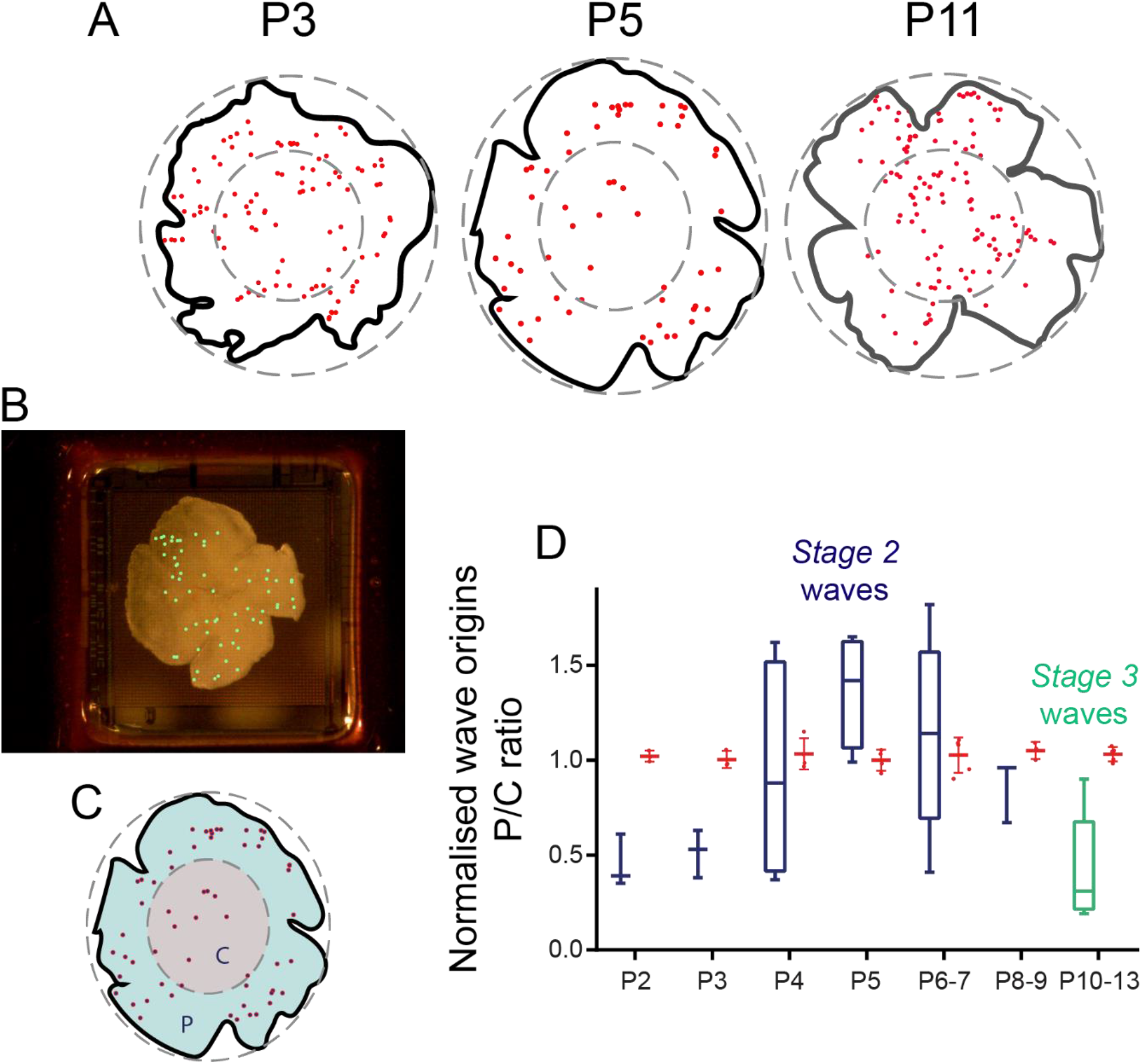
retinal waves and cluster cells. **A**: outlines of retinal wholemounts (black lines) photographed on the MEA immediately post recording (see **B** as well) overlaid with wave origins (red dots) detected during 30 minutes recording. Large grey dotted ellipses: encompass the whole retina. The smaller concentric ellipses (50% smaller than the large ones) indicate the central area. **B**: Photograph of a P4 retina on the MEA, taken at the end of the recording session. Wave origins (green dots) are overlaid on the photograph. **C**: Method to measure wave origins in center and periphery. P: periphery, C: center. Red dots: wave origins. **D**: boxplot illustrating the P/C ratios from P2-13. Wave origins expand from center to periphery between P2 and P5-6, similar to the clusters themselves (Figure 1C). Same boxplot conventions as for Figure 1C. Statistical analysis was not possible here due to the small numbers of values in each group (one ratio value per retina). Blue: Stage2 cholinergic waves; green: Stage3 glutamatergic waves; red: Monte Carlo randomized P/C ratio values.

To understand the possible functional involvement of these cellular clusters in wave generation, we used electrical imaging (Greschner et al, 2016; Zeck et al., 2011; Petrusca et al., 2007; Litke et al, 2004) (see Methods) to visualize wave-related electrical activity at high spatiotemporal resolution (Figure 4). Recordings were done on MEAs with either 81 μm or 42 μm electrode pitch, the latter providing higher spatial resolution. Although negative deflections smaller than the spike used for spike-triggered averaging (STA) were detected around most STA channels, presumably reflecting wave-related activity propagation (as in Channel 1782, Figure 4A and B), on some channels we could see conspicuous positive signals emerging simultaneously with the negative spike signal (Channel 1643, Figure 4A). These signals were significantly smaller (red asterisk) than the STA spikes (blue asterisk), but nevertheless easily distinct from baseline activity. Overall, when combining the activity footprint from all channels exhibiting such positive-negative “dipole” behavior (Figure 4C), we found that these areas form clusters in close proximity with wave origins (green dots in Figure 4C). However, when plotting all maximal projections, regardless of whether they have a positive deflection or not (Figure 4C, bottom row), the activity is more spread out over MEA channels, with less clear co-localization with the wave origins. The clustered layout of the dipole areas suggests that these signals may reflect activity originating from the transient cell clusters.

**Figure 4:**
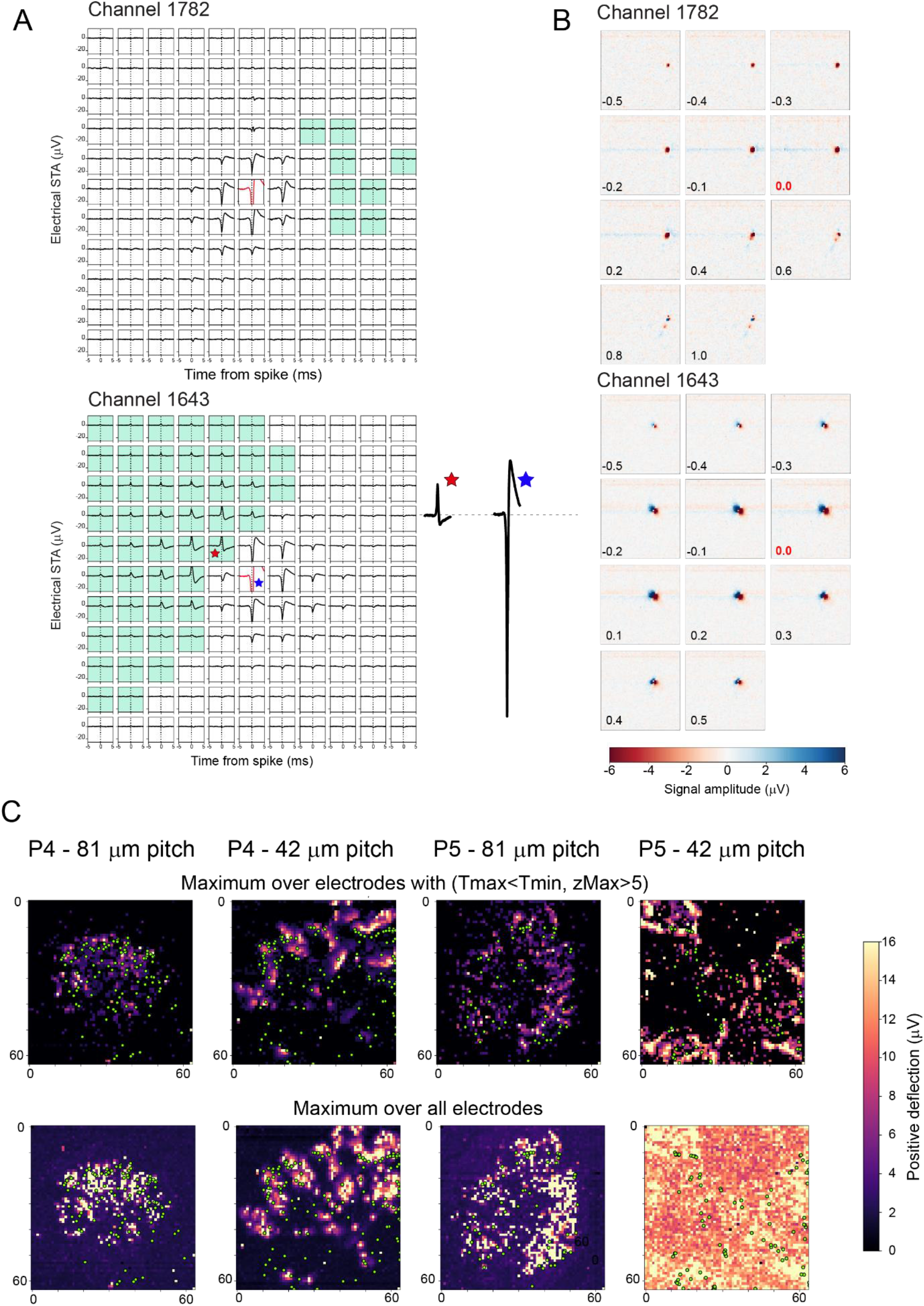
electrical imaging of retinal waves. **A**: STAs for two trigger channels showing signals averaged over - 5 to +5 ms relative to spikes on the trigger channel (in red) for 11×11 surrounding recording channels. Recording channels with dipole activity (with maximal positive deflections with z score>5 occurring before the maximal negative deflection, see Methods) are marked with the green mask. Channel 1643 has a marked area with dipole signals near the trigger channel. The single traces on the right side of the electrode grid show the full size of the spike on the trigger channel (blue asterisk) and maximal positive deflection (red asterisk), emphasizing the fact that the amplitude of the dipole signals is significantly smaller than spikes, suggesting that they may represent slow, graded potentials. (P4 retina, 60 min recording). **B**: Time-lapse images taken from movies of the averaged activity for both channels illustrated in **A**. The precise time of each image is indicated in the bottom left corner of each frame (in ms). Time 0.0 (time of the spikes used for STA) is indicated in red. For Channel 1643, clear dipole signals are seen from the earliest time frame (−0.5ms). Such signals are absent in Channel 1782. **C**: Maximal projections for signals with pre-STA spike signals with larger positive than negative deflections (top row) and for all maximal projections (bottom row) over the entire MEA. Maps are shown for two P4 and two P5 retinas, each with one example recorded on an array with 42 μm electrode pitch, and for another array with 81 μm pitch. Wave origins (green dots) are overlaid on the electrical signals.

## Discussion

Here we report for the first time that clusters of cells are transiently present in the mouse RGC layer during the period of Stage-2 cholinergic waves. They appear near the ONH at P2 and migrate to the periphery by P7 before disappearing at P10, coinciding with the switch to Stage-3 waves.

One plausible explanation that these cells have not been reported in previous studies is that they are sparse, and therefore may be easily missed if not visualized at pan-retinal level. Moreover, they auto-fluoresce, hence they could be mistaken for other cell types when using immunofluorescence because they give signals at most wavelengths (except in the UV range), even when not using any fluorescent antibodies (primary or secondary). Because of their auto-fluorescence, we have not been able to establish their cellular identity yet and other methods such as single-cell sequencing and non-fluorescence based immuno-labelling approaches will be necessary. The only thing we can claim with certainty at this point is that these cells are present only in the RGC layer, not in the INL. They populate the entire thickness of the GCL, and are often seen to contact SACs, suggesting that they may signal to these cells. They appear significantly larger than SACs and RGCs and they do not form a well-organized horizontal network spanning the entire retinal surface. Because of their location, they may represent a transient, specialized population of RGCs. However, we have not been able to detect any sign of apoptotic activity in these cells (using caspase3 immunolabelling, data not shown) whilst RGCs undergo massive programmed cell death during the same period (Young, 1984). It is unlikely that they are undifferentiated cells because they do not express the transcription factor *Olig2* (data not shown), widely expressed in retinal progenitor cells (Hafler et al, 2012).

Preliminary observations (not shown) from our lab suggest that they may die through microglial phagocytosis, a phenomenon that has been reported for neuron progenitors in the developing cortex (Cunningham et al., 2013). It is unlikely that these cell clusters physically move from the ONH to the periphery over such a short developmental period. It may be that they are born and differentiate in a center-to-periphery sequence, a well-known phenomenon during retinal development and at the same time, they are eliminated in gradually more eccentric distances from the ONH, creating the illusion of an eccentric migratory movement. We do not know why these cell are auto-fluorescent. It may be that the cells are initially not fluorescent, and become auto-fluorescent as a sign of stress that develops only when the cells are triggered to disappear, perhaps prompted by microglial contact.

Our anatomical and physiological observations suggest that these cells may be responsible for the generation of Stage-2 waves. Indeed, using an MEA that is large enough to cover the entire retina, we found that wave initiation points move in a non-random, center-to-periphery fashion over the same time frame and pattern as the clusters. Waves expand in size up to P6 (Maccione et al, 2014), which could be due to their initiation points becoming gradually more peripheral. Unfortunately, due to shrinkage and repetitive handling of fixed retinas, we have not yet been able to reliably compare the locations of cholinergic clusters with wave origin points in corresponding living retinas.

Electrical imaging analysis demonstrates the presence of activity clusters characterized by simultaneous positive and negative small signals in proximity with wave origins, suggesting that these clusters may represent activity related to the cholinergic RGCs. We propose that these cells act as hyper-excitable and hyper-connected hubs that trigger waves. Once generated, waves then travel across the retinal surface via the SAC network, as established in previous studies. Future studies are needed to reach a better understanding of the nature of the signals generated by these cells, and the extent and nature of functional connectivity they make with SACs and other RGCs.

Although these cells have never been reported in the retina, developing cortical areas exhibit a transient population of subplate neurons that are highly active and synaptically connected to other developing neurons (Luhmann et al, 2018). The presence of transient, electrically active neurons during early development is thus not a new concept, suggesting a universal mechanism mediating hyper-excitability in developing CNS networks during the critical period for brain wiring..

## Methods

### Animals

All experimental procedures were approved by the UK Home Office, Animals (Scientific procedures) Act 1986.

Experiments were performed on neonatal (P2-13) C57bl/6 mice. All animals were killed by cervical dislocation followed by enucleation.

### Immunostaining and imaging

For our anatomical studies, retinas were extracted from pups aged P2 (N=18 retinas), P3 (N=19), P4 (N=19), P5 (N=19), P6 (N=16), P7 (N=13), P8 (N=10), P9 (N=10), P10 (N=4) and P11 (N=4).

We have used the following antibodies:

### Primary antibodies

ChAT (AB144P, goat polyclonal, Merck Millipore).

VAChT (PA5-77386, rabbit polyclonal, ThermoFisher Scientific).

RBPMS (1830-RBPMS, rabbit polyclonal, Phosphosolutions).

### Secondary antibodies

Donkey anti rabbit Alexa 568 (A10042, Invitrogen).

Donkey anti goat Dylight 488 (SA5-10086, ThermoFisher Scientific).

### Retinal sections

Eyecups were prepared from mouse pups aged P2-P9, fixed for 45 min in 4% paraformaldehyde (PFA),), incubated in 30% sucrose in 0.1M phosphate buffer solution (PBS) for at least 12 hours, and then embedded in Optimal Cutting Temperature (OCT) cryo embedding compound and frozen at -20°C. Eyecups were sliced as 28µm thick sections using a cryostat (Model: OTF5000, Bright Instruments), washed with PBS to remove OCT, and incubated in blocking solution for 1 hour (5% secondary antibody host species serum with 0.5% Triton X-100 in PBS) prior to staining with antibodies.

Retinal sections were incubated with the primary antibody solution (0.5% Triton X-100 with VAChT (1:500 in PBS) and ChAT (1:500)) for 12 hours at 4°C. Sections were washed with PBS, followed by incubation with fluorescent secondary antibody solution (0.5% Triton X-100 with donkey anti rabbit Alexa 568 (1:500) and donkey anti goat Dylight 488 (1:500) in PBS) for 1 hour.

Finally, slices were washed with PBS and embedded with home-made OPTIClear refractive-index homogenisation solution. OPTIClear solution consists of 20% w/v N-methylglucamine, 25% w/v 2,2’-Thiodiethanol, 32% w/v Iohexol, pH 7-8. The solution is clear and colourless, with a refractive index of 1.47-1.48.

Sections were imaged using the Zeiss LSM 800 confocal microscope. Regions of interest were selected by focusing on clusters.

### Retinal wholemounts

Wholemount retinas were prepared from mouse pups aged P2-P11, flattened on nitrocellulose membrane filters and fixed for 45 min in 4% PFA. Retinas were then incubated in blocking solution (5% secondary antibody host species serum with 0.5% Triton X-100 in PBS) for 1 hour.

Primary antibodies: 0.5% Triton X-100 with RBPMS (1:500) and ChAT (1:500). Secondary antibodies: 0.5% Triton X-100 with donkey anti rabbit Alexa 568 (1:500) and donkey anti goat Dylight 488 (1:500).

Retinas were incubated with the primary antibody solution for 3 days at 4°C, then washed with PBS and incubated with the secondary antibody solution for 1 day at 4°C.

Finally, retinas were washed with PBS and embedded with OptiClear. Zeiss AxioImager with Apotome processing and the Zeiss LSM 800 confocal microscope were used to image the retinas.

High-resolution images of the RGC layer down to the INL were obtained by subdividing retinal wholemounts into adjacent smaller images that were subsequently stitched back together to view the entire retinal surface. Regions of interests were selected around the clusters.

To compensate for variability in retinal thickness, several focus points were set across the retinal surface in order to keep sharp focus on the desired cell layer. Each individual picture was then acquired in all color channels at 20x magnification, and with 10% overlap between neighboring areas. This overlap is used to correctly align and stitch together all pictures using the Zen Pro software (Zeiss). Z-stacks of images at 40x magnification were acquired at regions of interest to visualize cells in 3D. Z-stacks consisted of images taken every 1 µm from the RGC layer to below the INL.

To calculate the relative position of the cell clusters between the ONH and periphery, lines were traced and measured from the middle of the ONH to the middle of a cluster (D1) and then from the same point in the cluster to the periphery of the retina (D2). D1/D2 represents the relative position of the clusters. One-way ANOVA was used on all 233 ratio values for all eight groups. Tukey post-hoc test was used to identify significant changes in cluster positions between consecutive developmental days.

### Electrophysiology

#### MEA recordings

Retinas were isolated from mouse pups P2 (N=4 retinas), P3 (N=4), P4 (N=7), P5 (N=8), P6 (N=3), P7 (N=2), P8 (N=2), P9 (N=2), P10 (N=2), P11 (N=2), P12 (N=1), P13 (N=1). The isolated retina was placed, RGC layer facing down, onto the MEA and maintained stable by placing a small piece of polyester membrane filter (Sterlitech, Kent, WA, USA) on the retina followed by a home-made anchor. The retina was kept in constant darkness at 32°C with an in-line heater (Warner Instruments, Hamden, CT, USA) and continuously perfused using a peristaltic pump (∼1 ml/min) with artificial cerebrospinal fluid containing the following (in mM): 118 NaCl, 25 NaHCO_3_, 1 NaH_2_PO_4_, 3 KCl, 1 MgCl_2_, 2 CaCl_2_, and 10 glucose, equilibrated with 95% O_2_ and 5% CO_2_. Retinas were allowed to settle for 2 hours before recording, allowing sufficient time for spontaneous activity to reach steady-state levels.

High resolution extracellular recordings of spontaneous waves were performed as described in details in Maccione et al. (2014), using the BioCam4096 platform with APS MEA chips type HD-MEA Stimulo (3Brain GmbH, Switzerland), providing 4096 square microelectrodes of 21 µm x 21 µm in size on an active area of 5.12 × 5.12 mm, with an electrode pitch of 81 µm. Two P5 and one P4 datasets were acquired with the MEA chip HD-MEA Arena (2.67×2.67mm active area, electrode pitch 42 µm).

Raw signals were visualized and recorded at 7 kHz sampling rate with BrainWaveX (3Brain GmbH, Switzerland). Each dataset consisted of 30 min of continuous recording of retinal waves. The datasets used for electrical imaging were acquired at 17.855 kHz for 30 or 60 min.

In the BioCam4096, samples of MEA signal are acquired row-wise by the amplifier. Individual samples consist of 64 columns and 64 rows and often show a small but measurable bias across rows of ca. 2-4 µV (1-2 ADC units). While such bias is negligible for most applications, it does degrade the quality of electrical images. Therefore, to reduce the bias before electrical imaging, the median value of each row was independently calculated and subtracted.

Retinas were photographed on the MEA at the end of the recording session to ensure we document the precise orientation of the retina with respect to the array of electrodes (Figure 3B).

#### Data processing and analysis

Burst and wave detection was done in Matlab (Mathworks) as described in Maccione et al. (2014). We computed the coordinates of the wave origins (Maccione et al., 2014) and aligned them with a picture of the retina on the MEA (Figure 3B). Retinas were delimited by a peripheral annulus (P), surrounding a concentric central ellipse, 50% smaller (C) (Figure 3A and C). We counted wave origins (Figure 3C, red dots) localized in C (grey shading) and in the retinal area confined within P (green shading) and plotted the P/C ratio between P2-13 (Figure 3D).

Electrical images were computed independently for each electrode by averaging the electrical activity in the MEA surrounding the time of spikes in that electrode (spike-triggered average). First, spikes were detected independently at each electrode with the default detection parameters of BrainWaveX (3Brain GmbH, Switzerland) (Maccione et al., 2014). To ignore electrodes without good physical coupling to the retina, only active electrodes – those with any noteworthy activity – were analyzed. Active electrodes were defined as having a normalized spike count

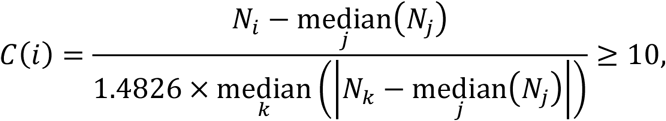

where *N* stands for the spike count from an electrode. The denominator of *C* robustly estimates the standard deviation of spike counts using the median absolute deviation from the median (Quiroga et al., 2004; Donoho & Johnstone, 1994).

The electrical imaging itself proceeded as follows. For each active electrode, snippets of raw MEA signals were taken from -5 to +5 ms of the detected spike times. Only spikes appearing concomitantly with retinal waves were considered (wave onset and offset were determined by (i) detecting bursts on each electrode separately, and (ii) grouping bursts into waves based on temporal overlap and proximity, see Maccione et al., 2014 for further details). For the sampling rate of 17.855 kHz, a snippet consisted of 64 rows, 64 columns, and 180 sample points. Averaging the snippets thus led to a 64×64×180 movie *A*_*i*_(*x, y, t*) representing the typical electrical activity in the temporal vicinity of spikes from electrode *i* (Figure 4B). To remove noisy electrodes, electrical images whose negative peak of the spike had a half-peak width of less than 0.3 ms were discarded from the remaining analysis. To visualise the electrical activity, movies *A*_*i*_(*x, y, t*) were reduced to a map of negative deflections 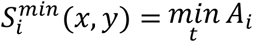, positive deflections 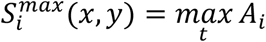, and the times 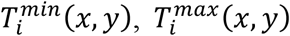 where such deflections occurred.

In some electrical images, a dipole activity was observed where positive deflections emerged simultaneously with the expected negative deflections of a spike. These positive lobes showed a strong positive deflection followed by a negative deflection (Figure 4A). To detect candidate regions with positive lobes, pixels (x,y) were selected when they had a significant positive deflection at a time 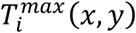 preceding the time of negative deflection 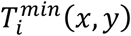 and not occurring later than 0.5 ms after the triggering spike. The latter condition helped to remove pixels exhibiting purely axonal propagation. Positive deflections were considered significant when their z-scored values

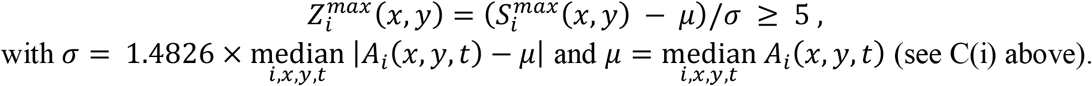

Defining a mask *W*_*i*_ (*x, y*) as unity for such selected pixels with positive deflections (green background in Figure 4A) and zero everywhere else, the maximum projection (Figure 4C) over selected regions was given by 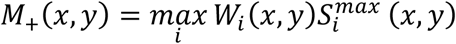. The maximum over all electrodes, regardless of positive lobes, was given by 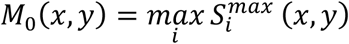.

## Acknowledgments

This work was supported by the Biotechnology and Biological Sciences Research Council (BBSRC, BH163322), Newcastle University Faculty of Medical Sciences and by the European Research Council (ERC) under the European Union’s Horizon 2020 research and innovation programme (grant agreement number 724822).

JdM, VK and ES designed the experiments; JdM and VK performed the experiments; JdM and ES analyzed the experimental data; FR and TG designed and performed the electrical imaging analysis; ES and FR wrote the manuscript with input from the other authors.

## Competing interests

We declare that we have no competing interests related to this submission.

## Notes

### Competing Interest Statement

The authors have declared no competing interest.

### Summary of Updates

In the original version we described the cluster cells as being a type of cholinergic retinal ganglion cell. In the meantime, we have discovered that these cells are auto-fluorescent, hence we cannot ascertain their identity by immuno-fluorescence because they give intrinsic fluorescent signals over a broad range of wavelengths. We have therefore removed the conclusion that the cluster cells are cholinergic ganglion cells and modified Figure 2 accordingly. We have also added some analysis of retinal waves, providing strong evidence that the wave origin locations are not random and follow a center-to-periphery pattern similar to the cell clusters during the first postnatal week.

